# Touch inhibits touch: sanshool-induced paradoxical tingling reveals perceptual interference between somatosensory submodalities

**DOI:** 10.1101/2020.11.21.391458

**Authors:** Antonio Cataldo, Nobuhiro Hagura, Yousef Hyder, Patrick Haggard

**Affiliations:** Institute of Cognitive Neuroscience, University College London, Alexandra House 17 Queen Square, London, WC1N 3AZ, UK; Institute of Philosophy, School of Advanced Study – University of London, Senate House, Malet Street, London, WC1E 7HU, UK; Cognition, Values and Behaviour, Ludwig Maximilian University, Gabelsbergerstraße 62, 80333 München, Germany; Center for Information and Neural Networks, National Institute of Information and Communications Technology, 1-4 Yamadaoka, Suita City, Osaka, 565-0871, Japan; Graduate School of Frontier Biosciences, Osaka University, Osaka, Japan

**Keywords:** Mechanoreceptor channel, SA mechanoreceptors, RA mechanoreceptors, Szechuan pepper, hydroxy-α-Sanshool, tactile perception

## Abstract

Human perception of touch is mediated by inputs from multiple channels. Classical theories postulate independent contributions of each channel to each tactile feature, with little or no interaction between channels. In contrast to this view, we show that inputs from two sub-modalities of mechanical input channels interact to determine tactile perception. The flutter-range vibration channel was activated anomalously using *hydroxy-α-sanshool*, a bioactive compound of Szechuan pepper, which chemically induces tingling sensations. We tested whether this tingling sensation on the lips was modulated by sustained mechanical pressure. Across four experiments, we show that sustained touch inhibits sanshool tingling sensations in a location-specific, pressure-level and time-dependent manner. Additional experiments ruled out mediation of nociceptive or affective (C-tactile) channels underlying this interaction. These results reveal novel inhibitory influence from steady-pressure onto flutter-range tactile perceptual channels, consistent with early-stage interactions between mechanoreceptor inputs within the somatosensory pathway.

## Introduction

The sense of touch involves neural processing of multiple features of cutaneous stimuli. Features extracted from stimuli to the skin are conveyed to the brain through distinct classes of afferent fibre [1,2]. Some fibres are tuned for specific spatiotemporal skin deformation patterns, and are considered mechanoreceptor channels, while others are tuned for thermal and noxious features [3,4]. These neurophysiological channels can also be studied psychophysically, because different qualities of sensation (flutter, high-frequency vibration, steady pressure, etc) are thought to be conveyed by each afferent class [1,2,5].

Although the characteristics of each perceptual channel have been explored, little is known about how the information from each channel interacts to provide an overall sense of touch. For example, inhibitory interaction between mechanical and pain/thermal channels has been well established [6,7], but it is still unclear whether similar inhibitory interactions occur between the different mechanoreceptor channels or ‘submodalities’. Classical accounts assume that each mechanoreceptor channel (RA, SA1, PC, SA2) carries *independent* information about specific tactile features [8,9], and that this independence is preserved in early cortical somatosensory processing [10–13]. The independence hypothesis has been recently challenged by neurophysiological studies of responses in single neurons. These studies suggested interaction between signals from different mechanoreceptor channels at spinal, thalamic and cortical levels [2,14,15]. However, to our knowledge, few psychophysical studies have investigated the implications of inter-channel interaction for perception, as opposed to neural coding.

Here, we show the first *human psychophysical* evidence that signals from different mechanical feature channels do indeed interact to determine tactile perception. Specifically, we show that perception of flutter-range mechanical vibration (mediated by a perceptual channel putatively corresponding to a rapidly adapting [RA] neurophysiological channel) is inhibited by concurrent activation of the perceptual channel for steady pressure (putatively corresponding to a slowly adapting [SA] channel). Thus, “touch inhibits touch”, in a manner similar to the established inhibitory interaction between mechanoreceptive and nociceptive channels (i.e. “touch inhibits pain”) [6,7].

Testing for interaction between perceptual channels might logically involve psychophysical tests of frequency-specific stimuli both alone, and in combination. However, delivering pure frequency-resolved stimuli to mechanoreceptors is difficult, because of the complex propagation of mechanical stimuli through the skin [16]. Here, we take an alternative approach that avoids the difficulties of delivering multi-channel mechanical stimuli, by *chemically* activating one target tactile feature channel, and then measuring the resulting percept in the presence or absence of additional mechanical stimulation to a second channel. In particular, we activated the perceptual flutter-range vibration channel (corresponding to a putative RA channel) using *hydroxyl-a-sanshool*, a bioactive compound of Szechuan pepper (hereafter sanshool), that produces localized tingling sensations with distinctive tactile qualities. Others have previously demonstrated that sanshool activates the light touch RA fibres [17–20], and we have previously shown that indeed, the perceptual flutter range tactile feature channel is activated by sanshool [21,22]. Here, we report the perceptual effects of first inducing sanshool-induced tingling, and then additionally applying controlled sustained pressure (corresponding to the putative SA channel input) to the same skin region. We used psychophysical methods to investigate how the intensity of sanshool-induced tingling sensations was modulated by the additional steady pressure input. This allowed us to assess the interaction between the two perceptual channels that are responsible for tactile steady pressure and tactile flutter features.

## Materials & Methods

### Participants

A total of 55 right-handed participants (age range: 18-38 years) volunteered in Experiments 1-5 (Experiment 1: 10 (two females), Experiment 2: 10 (5 females), Experiment 3: 9 (6 females), Experiment 4: 18 (13 females), Experiment 5: 8 (7 females). Fifty-one new participants (31 females) took part in Experiment 6, which was conducted in the context of a science and culture fair. All participants were naive regarding the experimental purpose and gave informed written consent. Experiments 1-5 were approved by University College London Research Ethics Committee. Experiment 6 was approved by the Research Ethics Committee of the School of Advanced Study, University of London. See Supplementary Material for the inclusion criteria of each experiment.

### Experiment 1

In Experiment 1 (n = 10), we tested whether the tingling sensation induced by sanshool (putative RA channel activation [17–23]) is modulated by application of sustained light pressure (putative SA channel activation [1,2,24]).

Tingling sensation was induced on the upper and lower lip vermilions by applying *hydroxyl-alpha-sanshool* (ZANTHALENE, 20% solution, Indena SPA., Milan, Italy) using a cotton swab (Figure 1A). This stimulation site was chosen because of its dense innervation of mechanoreceptors [25] and thin epidermis [26], which allows the chemical to reach the receptor effectively [21]. Participants sat on a chair, maintaining the upper and lower lip apart by biting a small section of straw between their canine teeth. Each trial started with a baseline period in which participants experienced the tingling sensation of sanshool all over the lips. Then, one of eight locations on the lips were manually stimulated by the experimenter with a probe (diameter: 14 mm, force: ~1.5 N) for 10 seconds. Touched locations included three positions each on the upper and lower lip vermillion and two positions above and below the vermillion border respectively (Figure 1A). Participants were instructed to always attend to the medial part of the lower lip (position 6 in Figure 1A), and to judge the intensity of tingling in this specific target location, while the sustained pressure probe contacted one of the eight locations on the lips. Thus, in the baseline condition before the application of the pressure probe, participants experienced the sanshool tingling sensation and were asked to memorise this baseline intensity. Next, the experimenter applied the mechanical stimulus probe, and the participants again rated the tingling sensation that they experienced, relative to the previous baseline, while the contact with the mechanical probe remained in steady contact with the skin. A rating of 10 indicated that the perceived tingling was at the same level as the intensity at baseline; a rating of 0 meant that the participant did not perceive any tingling sensation at all; ratings above 10 would indicate a higher tingling intensity than the baseline period. The rating was given 10 s after the mechanical probe had been applied, to minimise any transient mechanical effects. An inter-trial interval of a few seconds without mechanical stimulation was always included, to allow the tingling sensation to return. Return of the tingle was asked after every trial, and the next trial started only when this was confirmed by the participant. The experiment consisted of six blocks. Each block, consisted of 10 trials; 2 trials each on positions 3 and 6, and one for the remaining six positions (positions 1, 2, 4, 5, 7, 8). This was done to increase sensitivity for the conditions we thought more relevant to the interaction hypothesis. The order of locations for mechanical stimulation was randomised within each participant.

**Figure 1.**
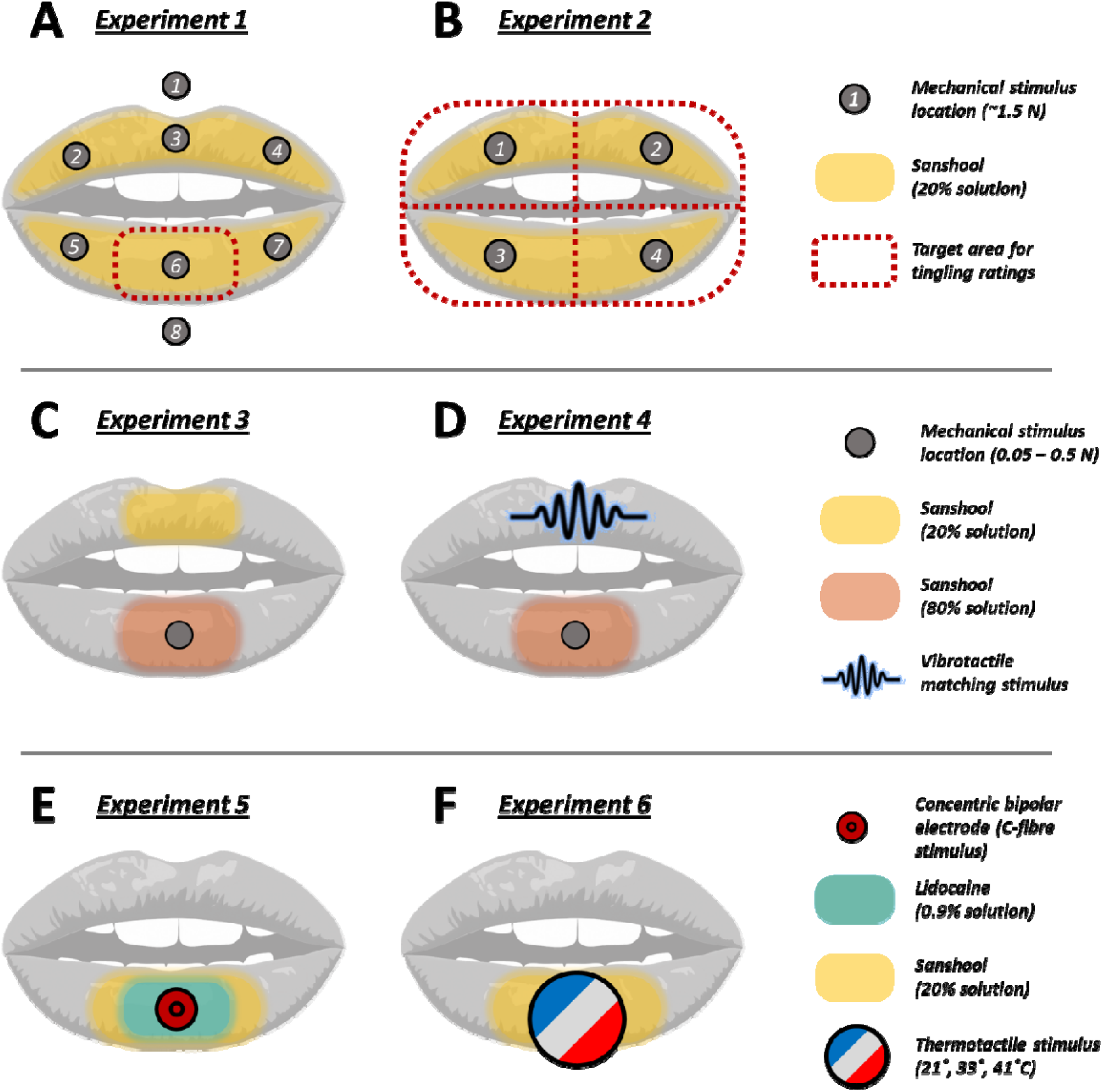
Experimental methods. **A-B:** In Experiments 1 and 2, participants (n = 10 in each experiment) experienced sanshool tingling all over the lips, while sustained touch (~1.5 N) was manually applied in different locations for 10 s. Participants reported the effect of touch location on sanshool tingling by rating the change in tingling intensity on the centre of the lower lip (**A:** Experiment 1) or all over the lips (**B:** Experiment 2). **C:** In Experiments 3, weaker and stronger sanshool solutions caused weaker and stronger tingling intensities on the upper and lower lips, respectively. Different levels of sustained force (0.05, 0.1625, 0.275, 0.375, and 0.5 N) were then applied to the lower lip by a closed-loop robotic device (see Figure 2). Participants (n = 8) reported which lip had the strongest tingling, as a function of sustained force. **D:** In Experiment 4, participants (n = 14) estimated the intensity of sanshool tingling on the lower lip by adjusting the amplitude of 50 Hz vibration applied to the upper lip until the intensities felt equal. Meanwhile, different levels of sustained force (0.05, 0.20 or 0.35 N) were applied on the lower lip. The adjustment was done at four different timings from the onset of the force (before pressure, and at 0, 5, and 10 s after pressure) (see Supplementary Video S1 at https://tinyurl.com/yyuoecqd for an example of a trial). **E:** In Experiment 5, the pain thresholds and tingling ratings of eight participants were measured during sanshool stimulation, both before and after topical application of Lidocaine (0.9%w/w). **F:** In Experiment 6, participants (n = 51) rated the intensity of sanshool tingling (20%) during three levels of thermotactile stimulation (21, 33, and 41° C).

### Experiment 2

Experiment 2 aimed to replicate and generalise the results of Experiment 1. The procedure was largely similar to Experiment 1. To make sure that the effect obtained in Experiment 1 was not due to sustained spatial attention to a single target location, participants experienced sanshool tingling all over the lips, and sustained pressure was applied to one of four quadrants (Figure 1B) randomly chosen on each trial. This time, instead of only rating the sanshool tingling at a single, fixed location, participants gave separate ratings of tingling intensity for all four lip quadrants, with the order of prompting being randomised. Participants completed six blocks. In each block, sustained touch was applied once to each location (16 ratings).

### Experiment 3

Experiment 3 investigated whether sanshool tingling is modulated by different contact force levels. Given that SA receptor firing is proportional to contact force, [1,2,24], any neural interaction between the putative SA channel and sanshool-evoked tingling (putative RA channel) sensation should produce attenuation of sanshool tingling proportional to contact force. We tested this hypothesis with a novel psychophysical method involving comparing the intensity of two tingle sensations.

First, we arranged a situation where tingling intensity was higher for the lower lip than the upper lip, by applying 80% and 20% concentration sanshool solutions to the lower and upper lip respectively (Figure 1C). Participants rested on a chinrest with their lips kept apart (Figure 2). Prior to the main experiment, we confirmed that the stronger solution level of sanshool (lower lip) induced stronger intensity of tingling sensation compared to the weak solution (upper lip) (Supplementary Figure S2). Next, the medial part of the lower lip, which experienced the stronger tingling sensation, was stimulated with different contact forces (0.05, 0.1625, 0.275, 0.375, and 0.5 N). Forces were applied by a closed-loop system comprising a linear actuator (ZABER, XYZ Series, Vancouver, Canada) and a force gauge (Mecmesin, PFI-200N GEB, Slinfold, UK) (Figure 2), which continuously maintained the desired pressure level. A cotton bud (diameter 4.5 mm) was placed between the force sensor and the lip. Participants performed a two-alternative forced choice comparison task to indicate whether the upper or the lower lip experienced the more intense tingling sensation. In each trial, one of five different levels of force were applied to the lower lip. One second after the onset of steady pressure, an auditory tone signalled that participants should judge whether the lower or the upper lip currently had the higher intensity of tingling sensation. Participants performed three blocks, each consisting of ten repetitions of the five contact forces, in random order, giving 150 trials in total.

**Figure 2.**
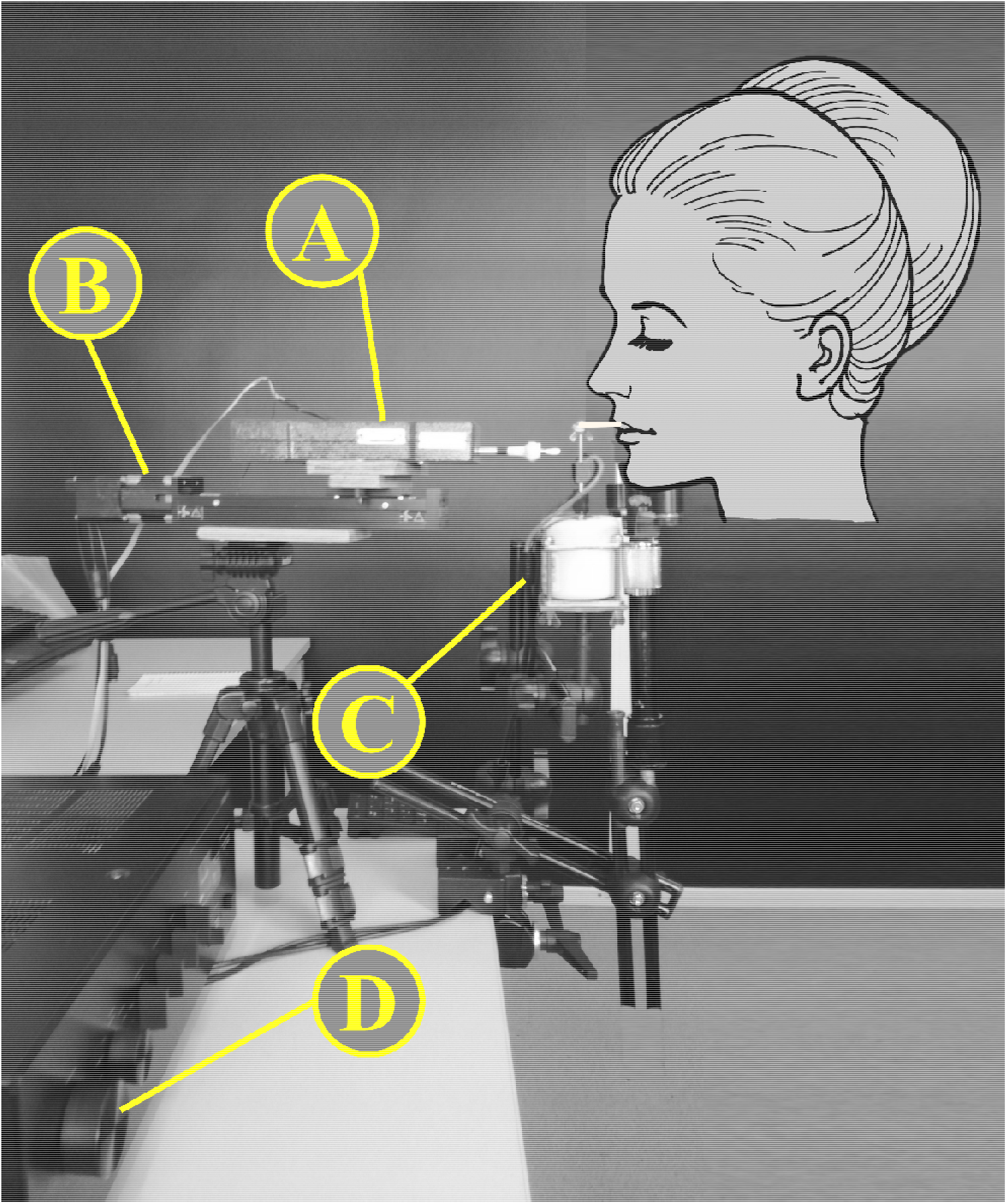
Experimental setup. In Experiments 3 and 4, sustained touch was applied to the lower lip by a mechanical probe, which exerted a target level of contact force via a motor (**B**) controlled in a closed-loop arrangement using a strain-gauge force sensor (**A**). In Experiment 4, 50 Hz mechanical vibration was applied to the upper lip by a vibrator (**C**). Participants could adjust the vibration amplitude using the gain knob of an amplifier (**D**). See Supplementary Video S1 at https://tinyurl.com/yyuoecqd for a video of the setup and an example trial of Experiment 4.

### Experiment 4

Experiment 4 tested how tingle intensity varied according to the time course of a sustained pressure stimulus. The discharge rate of SA neurons in response to static touch decreases gradually over time, dropping to 30% of the initial firing after 10 seconds [24]. Therefore, if the activation of the pressure (SA) channel drives suppression of the tingling sensation, some time-dependent recovery of tingling sensation should occur.

The setup was similar to Experiment 3 (Figure 1D). However, sanshool (80% solution) was applied on the lower lip only, while the upper lip rested on the probe of a vibro-tactile shaker (BRÜEL & KJÆR, LDS V101, Nærum, Denmark) (Figure 2). In each trial, participants first estimated the intensity of sanshool tingling on the lower lip by adjusting the amplitude of 50 Hz vibration [21] applied to the upper lip until the two intensities felt similar. Amplitude adjustments were made by the participant using the volume setting of an electronic amplifier. The point of perceptual equivalence between mechanical vibration and sanshool-evoked tingle was indicated by pressing a key. Next, one of three different force levels (0.05, 0.20 or 0.35 N) was applied to the lower lip (Figure 1D). An auditory signal was delivered when the closed-loop system had achieved a steady force at the target level. Participants were instructed to note the intensity of the tingling sensation on the lower lip at the time of the beep, and to adjust the amplitude of mechanical vibration to the upper lip until it had a perceptually equivalent intensity. They were instructed to make the adjustment as accurately as possible, while taking no longer than 5 s. Their mean reaction time was 2.30 s (SD 0.62 s). Two further beeps sounded 5 and 10 s after the initial application of sustained force contact, requiring two further matching attempts. Thus, four successive estimations were collected in each trial, one before and three after the pressure application. For a video of the setup and an example trial of Experiment 4, see Supplementary Video S1 at https://tinyurl.com/yyuoecqd. The experiment consisted of three blocks, with each block consisting of ten repetitions of the three force levels (30 trials in total). The order of the forces was randomised within each participant.

### Experiment 5

Experiment 5 investigated whether sanshool-induced tingling might reflect the activity of the nociceptive channels induced by the activation of nociceptive small-diameter C-fibres. Animal studies show that sanshool activates both large-diameter myelinated (Aβ) neurons, as well as small-diameter unmyelinated C-fibres [17,19,20]. Although nociceptive Aβ neurons have also been recently described in humans [27], it is not clear whether these fibres are also activated by sanshool. We reasoned that attenuation of sanshool tingling by steady pressure could only be considered a touch-touch interaction if the tingling sensation could be attributed to a perceptual channel for touch, rather than a nociceptive channel. Therefore, to rule out the contribution of nociceptive C-fibres to sanshool tingling, we measured both pain thresholds and perceived intensity of tingling before and after blocking the activity of small fibres through lidocaine [28,29]. If the tingling is unrelated to the C-fibre activation, the sensitivity to pain would be affected by lidocaine, but the intensity of the tingling sensation would not.

Sanshool (20% solution) was applied on the lower lip. Pain thresholds for electro-tactile stimuli delivered on the lips were obtained using 4 mm diameter concentric bipolar electrode connected to a constant current stimulator (Digitimer, Ltd., DS7, Welwyn Garden City, United Kingdom) (Figure 1E). This type of electrode is known to preferentially activate nociceptive fibres at low intensities [30]. Small diameter C-fibres were blocked through a 0.9%w/w lidocaine hydrochloride solution (Boots UK Ltd, ANBESOL liquid, Nottingham, United Kingdom), a non-prescription topical anaesthetic widely used for relief of orofacial pain. Lidocaine is thought to affect predominantly Nav1.7 channels [29] and to preferentially block nociceptive fibres [28]. While lidocaine also blocks large fibres [28,31], its anaesthetic effects on different submodalities display a distinct temporal gradient [31]. In particular, pinprick and thermal sensations that characterise nociceptive and thermoceptive activities are impaired within 10 minutes of application. Conversely, tactile, proprioceptive, and motor sensations associated with larger fibres activity are affected only after longer intervals after administration (~15 minutes) [31]. Thus, to ensure an effective block of smaller fibres only, tests were performed within 10 minutes of lidocaine application.

In a 2 (time: pre-block, post-block) x 2 (sensation judged: tingling, pain) within-subject design, pain thresholds and tingling intensity ratings were measured before and after administration of lidocaine. Participants performed two sessions. In session A, participants rated the intensity of sanshool tingling on the lower lip (from 0: “no tingling at all”, to 10: “the strongest tingling sensation imaginable”). Session B took place at least one hour after session A, when participants confirmed that the tingling sensation had completely disappeared [32]. First, participants’ pain thresholds were estimated using a staircase procedure [33] (see Supplementary Material for details), then lidocaine was applied on a 2 x 1 cm area in the centre of the lower lip. Pain thresholds were estimated again after three minutes. Immediately after the second pain threshold estimate, the same area of the lip was painted with sanshool, and participants rated the intensity of tingling, using the same scale used in session A.

### Experiment 6

Experiment 6 investigated whether sanshool tingle might reflect activation of C-tactile afferents. Small-diameter C-fibres responsive to tactile, but not to nociceptive stimuli have been characterised in detail by many animal [3,34] and human studies [35,36]. Although C-tactile fibres are commonly found in hairy, but not glabrous skin [36], there is electrophysiological evidence of the existence of C low-threshold mechanoreceptors in the glabrous skin of rat hind paw [37], and psychophysical evidence, using nerve blocks, of a C low-threshold input from the glabrous skin of human hand [38,39]. While we are not aware of reports of C-tactile fibres innervating the skin of the lips, this may simply reflect previous sampling, and the possibility cannot be excluded. C-tactile fibres generally respond preferentially to tactile stimuli moving at intermediate velocities [35,36]. Tactile motion tuning cannot readily be assessed with a chemical stimulus like sanshool. Importantly, however, C-tactile fibres also show preference for neutral, skin temperature (32 °C) stimuli, rather than warm (40 °C) or cold (18 °C) stimuli [35]. Thus, if the sanshool tingle is mediated by a C-tactile channel, the perceived intensity of sanshool tingling should be maximal at neutral temperatures and reduced during cold or warm thermal stimulation, producing an inverted U-shape.

The perceived intensity of sanshool-induced tingling on the lower lip (20% solution) was assessed during three different thermo-tactile conditions: cold (21 °C), neutral (33 °C), and warm touch (41 °C) (Figure 1F). As a baseline condition, tingling intensity without any stimulation was also measured using the same scale used in Experiment 5. Thermo-tactile stimuli were delivered using a 13 mm diameter Peltier thermode (Physitemp Instruments Inc, NTE-2A, New Jersey, USA). Each trial started with a 10 s countdown to allow the thermode to reach the intended temperature. Then participants applied their lower lip against the thermode probe. Participants were asked to move their head to approach the probe of the Peltier device in each trial until their lower lip contacted it, and then maintain a posture that applied gentle touch for 4 s. The experimenter monitored that attendees kept their lip in stable contact with the stimulator for the entire duration of the stimulation. Four seconds after the initial contact, participants were prompted to rate the intensity of the tingle. After the rating, participants withdrew their lip from the thermode. Each thermo-tactile condition was repeated four times (12 trials in total). The order of thermal conditions was randomised across participants.

## Results

### Experiment 1: Sustained light-touch (putative SA input) inhibits Sanshool-induced tingling (putative RA input)

When the probe was applied at the judged target position (always the centre of the lower lip), tingling intensity was dramatically reduced (to a mean 24.7% ± SD 34.0 of the perceived intensity at baseline before the probe was applied) (Figure 3A). A one-sample t-test was used to compare the perceived intensity of tingle when the probe was present, to the null mean value of 10 which was defined in our rating scale as the perceived intensity at baseline). The result showed a significant reduction (t(9) = 7.00, *p* < 0.001, dz = 2.21. All p-values are Bonferroni-corrected for 8 positions).

**Figure 3.**
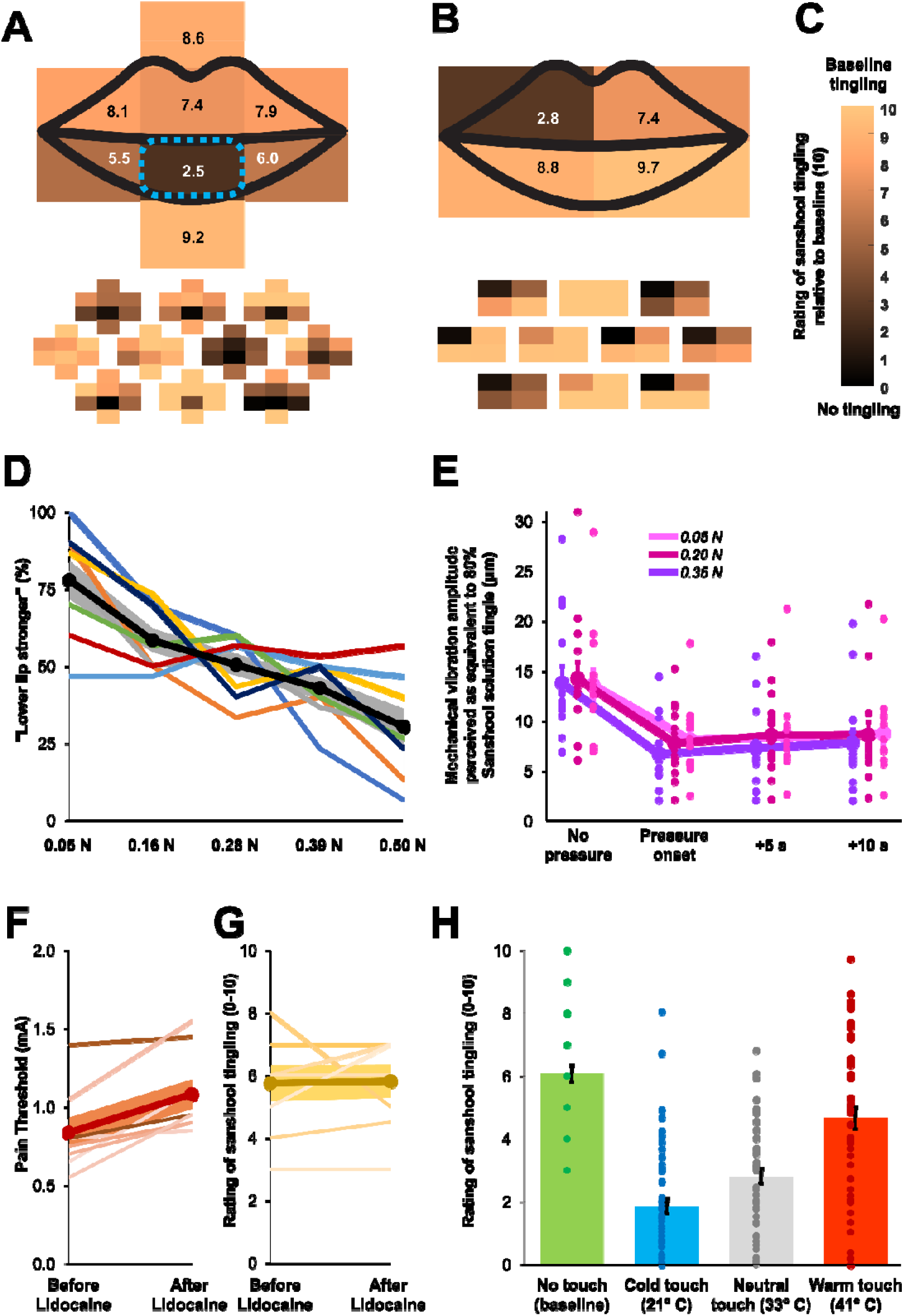
Results. **A-C**: Mean (top) and individual (bottom; n = 10 in each experiment) perceived intensity of sanshool tingling as a function of touch location in Experiments 1 (**A**) and 2 (**B**). Colour indicates the perceived intensity relative to the baseline period (**C**) (darker colours indicate lower ratings). In Experiment 1 (**A**), the perceived intensity of sanshool tingling at the target location (centre of lower lip) dropped significantly when sites on the lower lip were touched by the probe. In Experiment 2 (**B**), touch was applied to each of four quadrants in random order. Each time touch was applied, participants gave separate tingling ratings for each quadrant, after being prompted in random order. Data from all four touch conditions were realigned to the left upper lip position to express the spatial relation between the location where tingling intensity was judged and the location where sustained touch was applied. Significant intensity reduction was observed at the location where the touch was applied. **D:** In Experiment 3 (n = 8), the probability of judging the lower lip tingling intensity as stronger than the upper lip decreased as force level increased. The black line represents the sample average, the grey shading represents the SEM, and coloured lines represent individual data. **E:** In Experiment 4 (n = 14), the estimated vibration amplitude of sanshool tingling decreased as a static force increased. Moreover, the intensity of tingling significantly recovered as time elapsed. Error bars indicate the SEM across participants and coloured dots indicate individual data. **F-G:** Pain thresholds and ratings of sanshool tingling intensity before and after administration of lidocaine in Experiment 5 (n = 8). As expected, lidocaine induced a significant increase in participants’ pain thresholds (**F**). In contrast, the perceived intensity of sanshool-induced tingling was not affected by lidocaine administration (**G**). Dark lines represent the sample average, shadings represent the SEM, and coloured lines represent individual data. **H:** Participants’ ratings of sanshool-induced tingling during three thermo-tactile conditions in Experiment 6 (n = 51). The tingling intensity was linearly modulated by cold (21 °C), neutral (33 °C) and warm (41 °C) stimuli. Error bars indicate SEM across participants and coloured dots represent individual data.

The tingling sensation at the target position was not affected by pressure on the upper lip or off the lips (all *p* > 0.25, Bonferroni corrected). However, a significant reduction in tingling intensity relative to baseline was found when pressure was applied to the two lower lip locations adjacent to the judged target location (left side: t(9) = 4.28, *p* = 0.016 Bonferroni corrected, dz = 1.35; right side: t(9) = 4.25, *p* = 0.017 Bonferroni corrected, dz = 1.34). A repeated measures ANOVA showed a clear spatial gradient on the lower, but not the upper lip (see Supplementary Material).

Thus, sustained touch produced a robust inhibition of tingling sensation at the location where the tingling intensity was judged and at adjacent locations.

### Experiment 2: Inhibition of sanshool tingling sensation is spatially graded

For the quadrant where sustained touch was applied, we replicated the results of Experiment 1, finding robust reduction of tingling under pressure relative to the baseline (mean rating; 28.3% ± SD 36.8 of the baseline intensity) (see Supplementary Figure S1). We re-aligned the rating data of each remaining quadrant relative to the quadrant where the sustained touch was applied (Figure 3B). We could thus compare the effect on tingling of delivering sustained touch to either the same lip as the location where the tingling rating was judged, or the other lip, and likewise for sustained touch on the same side of the midline as the rated location, or the opposite side. The realigned data showed significant reduction of the tingling rating from the baseline at the quadrant where the sustained touch was applied (t(9) = 6.17, *p* < 0.001 Bonferroni corrected for four comparisons, dz = 1.95), and also at the other quadrant on the same lip (t(9) = 2.56, *p* = 0.045 corrected, dz = 1.00) (Figure 3B).

Next, we directly compared the tingle ratings across different locations in respect to the probe (realigned data). A 2 (lip; same or different to the probe) x 2 (side; same or different to the probe) repeated measures ANOVA revealed significant main effect both for the factor of the lip (*p* = 0.003, η_p_^2^ = 0.655) and the side of the probe (*p* < 0.001, η_p_^2^ = 0.842), and also an interaction effect (*p* = 0.001, η_p_^2^ = 0.692). In the planned comparisons, for the touched lip, the tingle ratings for the touched quadrant was significantly more inhibited compared to the untouched quadrant (t(9) = 6.30, *p* < 0.001, dz = 1.99). Interestingly, on the untouched lip also, the quadrant on the same side as the touch again had lower ratings than the other side (t(9) = 1.58, *p* = 0.025, dz = 0.50). This implies that the inhibition of the tingling depends on the spatial distance between the location where tingling is judged and the location of sustained touch, both within and across lips. Since the lips did not touch during the experiment (see Methods) this rules out mechanical propagation of sustained pressure as the cause of altered RA mechanoreceptor transduction. Instead, the interaction appears to occur at some neural processing level where afferents from the two mechanoreceptors are integrated in a spatially organised manner.

### Experiment 3: Sanshool tingling is parametrically inhibited as a function of contact force

We first checked that sanshool concentration influenced tingling intensity. As expected, participants reported significantly higher intensity for the 80% concentration on the lower lip (average rating: 6.6 ± SD 1.55) compared to the 20% concentration on the upper lip (average rating: 3.2 ± SD 1.06) (t(7) = 6.94, *p* < 0.001, dz = 2.45) (Supplementary Figure S2).

The probability of participants reporting a stronger sensation on the lower lip reduced progressively, and approximately linearly, as the force on the lower lip increased (F(1.6,11.2) = 12.09; *p* = 0.002; η_p_^2^ = 0.63) (Figure 3D). The suppressive effect of pressure on tingling intensity was confirmed by linear trend analysis (F(1,7) = 15.43; *p* = 0.006; η_p_^2^ = 0.69). Thus, RA activation induced by sanshool is parametrically modulated by the signal strength of the SA input.

### Experiment 4: Quantifying the relation between sustained force and sanshool– tingling sensation across time

After initial inspection of the data, we found that the distribution of the vibration amplitude matches deviated significantly from the normal distribution (see Supplementary Table S7). The statistical analysis was therefore conducted after log-transforming the data. However, to maintain the data in interpretable scale, we report and show the means and the standard errors in the original units (μm). The initial perceived tingling on the lower lip without pressure was matched by, on average, 13.9 μm (± SD 5.9) peak-to-peak amplitude of a 50 Hz vibration on the upper lip. Sustained contact force of 0.05 N on the lower lip reduced the tingling to a level that was now matched by 8.4 μm (± SD 4.6) of vibration amplitude. Contact forces of 0.20 N and 0.35 N were matched by 8.2 μm (± SD 4.1) and 7.6 μm (± SD 3.4) vibration amplitudes respectively (Figure 3E). For each contact force level, the perceived intensity of tingling was significantly reduced at all time points (pressure onset, +5 s, and +10 s after pressure onset) compared to the initial baseline period without any pressure contact (*p* < 0.05 corrected for all 9 comparisons), replicating the result of Experiments 1-3.

We specifically wanted to investigate whether the reduction of tingling sensation would change with the force level applied, and whether that reduction would change as a function of time from onset of probe contact. A 3 (force: 0.05, 0.20, and 0.35 N) x 3 (time: force onset, +5 s, and +10 s after force onset) repeated measures ANOVA on the vibration amplitude showed significant main effect for both factors of contact force level (F(2,26) = 4.32; *p* = 0.024; η_p_^2^ = 0.249) and time since contact (F(2,26) = 4.92; *p* = 0.015; η_p_^2^ = 0.275), but no significant interaction effect (F(4,52) = 0.27; *p* = 0.897) (Figure 3E).

We used Fisher’s LSD methods to identify conditions that differed significantly. For the force level factor, estimated tingling amplitude was significantly reduced in the highest (0.35 N) compared to the middle force level (0.20 N) (t(13) = 3.54, *p* = 0.004, dz = 0.94). Comparison with the lowest force level (0.05 N) showed a similar trend (t(13) = 2.15, *p* = 0.051, dz = 0.57) (Figure 3E). Therefore, the intensity of tingling was suppressed in a force-dependent fashion, as expected from Experiment 3. We investigated the effect of time in the same way. The perceived tingle intensity recovered as time elapsed (onset vs. +5 s: t(13) = 2.82, *p* = 0.014, dz = 0.75; onset vs. +10 s: t(13) = 2.38, *p* = 0.033, dz = 0.63) (Figure 3E). Since activity of SA neurons gradually reduces over time due to the adaptation to sustained pressure input [24], this modest time-dependent recovery of tingling sensation is consistent with the hypothesis that activity of SA neurons underlies suppression of RA mediated sanshool-tingling.

### Experiment 5: Sanshool-evoked tingling is not mediated by small-diameter unmyelinated C-nociceptive fibres

Bayesian statistics [40,41] were used as the experiment was designed to test the null hypothesis (i.e., lidocaine does not reduce sanshool tingle). First, we ran a one-tailed Bayesian t-test to confirm that lidocaine effectively blocked small C-nociceptive fibres. As expected, participants’ pain thresholds were significantly higher after lidocaine gel administration to the lips (mean 1.08 mA ± SD 0.27), compared with pre-administration (mean 0.84 mA ± SD 0.27) (BF_10_ = 18.26; error % < 0.001) (Figure 3F). We then tested our null hypothesis that lidocaine administration would not affect sanshool-induced tingling. As the alternative hypothesis (i.e. lidocaine reduces tingling) was unidirectional, a one-tailed test was used. The Bayesian analysis showed that the data were more likely under the null than under the alternative hypothesis (BF_01_ = 3.239; error % = 0.009). Tingling ratings were statistically identical before (mean 5.78 ± SD 1.7 arbitrary units) and after (mean 5.81 ± SD 1.5 arbitrary units) lidocaine administration (Figure 3G).

Thus, despite effective block of nociceptive afference by lidocaine, sanshool-evoked tingle remained unaltered, suggesting the small fibre C-nociceptors are not the main contributor on tingling sensation. Therefore, tactile gating of tingle presumably involves a different mechanism to the familiar “gate control” of pain by touch.

### Experiment 6: Affective touch channel activation does not mediate sanshool-evoked tingling

First, we confirmed that sanshool tingling is suppressed by the sustained touch, as shown in Experiments 1-4. As expected, sustained touch at neutral temperature significantly decreased (53.8%) the sanshool tingling intensity (mean rating 2.81 ± SD 1.7) compared with no mechanical touch (mean rating 6.08 ± SD 1.9) (t(50) = 9.01, *p* < 0.0001, dz = 1.26). Next, we compared ratings of tingling intensity under different temperature conditions. Cold touch produced the lowest ratings (1.85 ± SD 1.7), and warm touch the highest (4.67 ± SD 2.6) (Figure 3H). Therefore, the suppression effect decreased as the temperature of the stimulus increases. We accordingly found a significant main effect of temperature conditions in a oneway repeated measures ANOVA on participants’ intensity ratings of tingling during the three thermo-tactile conditions (F(1.5, 77.2) = 43.34; *p* < 0.0001; η_p_^2^ = 0.464). All pairwise comparisons were significant (*p* ≤ 0.002 in each case; Bonferroni-corrected). This suppression pattern clearly difìers from the inverted U-shape that would be expected from C-tactile fibre thermal sensitivity. On the other hand, the linear relation to temperature is consistent with the known thermal modulation of SA fibres, which respond more at lower temperatures (see Discussion).

## Discussion

Somatosensory perception involves integration of multiple features that reach the brain through different afferent channels. A central question is therefore whether and how inputs from these different channels interact with each other [2,14,42]. Classical theory suggested that specific frequency-selective channels, associated with specific receptors and afferent fibre types, were processed independently at least until early sensory cortex [10]. While some neuronal studies have begun to challenge the classical view of independent frequency channels [2,14,15,43], our study constitutes the first *perceptual* evidence for interactions between somatosensory submodalities. Using a novel approach involving anomalous chemical stimulation of mechanoreceptor channels, we show strong inhibitory interactions between distinct perceptual channels encoding different preferred frequencies. Specifically, we show that the tingling sensation associated with the flutter-range vibratory channel (putative RA channel) is inhibited by the input of sustained pressure (putative SA channel). We further showed that this inhibitory interaction is spatially selective and proportional to the activation of the pressure channel.

In the current study, we investigated perceptual channels based on psychophysically-defined characteristics. These methods identify perceptual channels by threshold differences across different stimulus frequencies, and by observing perceptual modulations due to adaptation and masking [44]. Although the peripheral (receptor/afferent fibre) basis of tactile feature processing have been extensively studied by neurophysiologists, we still do not know the precise details of the mapping between channels defined by peripheral physiology, and the perceptual channels defined by psychophysics. Nevertheless, the principle of studying principles of CNS organisation based on psychophysically-defined perceptual channels has been well established, for example in the visual system [45]. By analogy to visual psychophysics studies, we believe that the tactile feature processing system can also be usefully investigated by studying interaction between perceptual channels.

*Hydroxy-α-Sanshool* has been shown to activate the rapidly-adapting light-touch fibres in rats [17,20]. Using both adaptation [21] and masking paradigms [22], previous studies have demonstrated that a flutter-range vibration channel (putative RA channel) activation is responsible for the sanshool-induced tingling sensation. First, the perceived sanshool-induced tingling frequency on the lips is reduced by adapting the RA channel using prolonged mechanical vibration [21], paralleling the reduction in perceived frequency of mechanical vibration by similar adaptation procedures. Second, application of sanshool on the skin impairs detection of 30 Hz mechanical vibration (RA channel dominant frequency) but does not affect detection of 240 Hz (PC channel dominant frequency) or 1 Hz (SA channel dominant frequency) mechanical vibration [22], demonstrating that sanshool can selectively affect the putative RA channel. Finally, microstimuluation studies confirm the strong link between RA activation and flutter-range vibration sensations [46]. Thus, although we could not directly measure RA afferent responses to sanshool, we may nevertheless study the perceptually-defined channel underlying the sanshool tingling sensation, while identifying this putatively as an RA channel. Future microneurographic studies could potentially provide stronger evidence about the physiological afferents responsive to sanshool, including selectivity for particular afferent types.

In the meantime, psychophysical techniques can go some way to investigating whether other non-mechanical, temperature/pain related channels might also contribute to sanshool tingling sensations. While animal studies have shown that sanshool activates small fibres [17,20] as well as RA fibres, our perceptual Experiments 5 and 6 suggested that C-nociceptive and C-tactile fibres did not contribute to the tingling sensation. First, perceived intensity of sanshool-evoked tingling was unaffected by topical lidocaine anaesthetics that preferentially block small fibres [28,29]. Second, perceived intensity of the tingling sensation showed a linear and monotonic increase as a function of stimulation temperature, in clear contrast to the inverted-U shape that characterises thermal modulation of C-tactile firing [35]. However the thermal sensitivity of C-tactile fibres remains controversial, since some studies found C-tactile responses to cooling of the skin [47,48], rather than the inverted-U shape [35]. However, our observation of enhancement of tingling by warmth is incompatible with both of these reported patterns of C-tactile thermal modulation, yet is compatible with the reported thermal modulation of RA firing [49,50]. Our perceptual findings also agree with evidence from microneurography [51] and clinical neuropathies [52,53], which both consistently identify tingling paraesthesia sensations with activation of large-diameter afferents. In contrast, activation of small-diameter afferents generally elicits low, dull, painful sensations.

We found that sustained light touch attenuated sanshool tingle, and we propose that this reflects an interaction between the corresponding perceptual channels. Our experimental design successfully controlled for several alternative possible explanations of touch-induced suppression of tingling. First, we ruled out the possibility that sustained touch may have attracted attention to mechanical stimulus, either distracting attention away from the tingling sensation, or masking sanshool-induced activity in the same channel [54,55]. Explanations based on distraction cannot readily explain why suppression of tingle was location-dependent, with stronger suppression of tingling at the location of touch compared to remote from it. Alternative explanations based on masking would require the steady pressure stimulus to activate the same perceptual channel as sanshool, i.e., the putative RA channel. RA afferents typically respond at the onset of a steady pressure, but lack a sustained response [5,56]. Intra-channel masking theories would therefore predict transient suppression of tingle sensation at the onset of steady pressure, with rapid rebound of tingle during continued tactile contact. Yet, in Experiments 1, 2, and 4 we found significant touch-induced attenuation of sanshool tingle after 10 s of continuous touch, suggesting that an RA contribution to the attenuation of tingle is unlikely. Moreover, in Experiments 3 and 4 we found that pressure-induced attenuation of sanshool-evoked tingling increases linearly with the indentation. Linear increase in firing rate with indentation is a characteristic marker of SA fibres [24,57], but is absent in RA fibres, which are instead mostly affected by indentation velocity [56]. Thus, overall, the effects of steady pressure on sanshool-evoked tingling are consistent with steady pressure conveyed by a putative SA pathway, influencing sanshool-induced activity in a putative RA pathway.

Second, could our results reflect some perceptual filtering mechanism? For example, sustained pressure evokes a familiar and “meaningful” sensation, while the unnatural pattern of RA fibre activation by sanshool might be interpreted by the brain as a strange paraesthesia-like sensation. The brain might then prioritise interpretable touch signals, and filtering out the sanshool-induced signals. This top-down filtering account seems unlikely for at least two reasons. First, the account presupposes that sanshool-induced tingling is an ambiguous and unfamiliar sensation which the brain effectively discards. In fact, our participants could quickly and easily relate the tingling sensation to previous experiences, either spontaneously, or when prompted during debriefing. Common reports referred to: Szechuan cuisine, pins and needles, insects crawling on the lips, or “embouchure collapse” during brass playing. Sanshool-induced tingling is thus a clear, reportable and graded sensation, consistent with previous reports [21]. Second, our results clearly showed modulations of sanshool tingling that seem unrelated to familiarity and interpretability. For example, Experiment 6 showed that temperature of tactile stimulation systematically influenced sanshool tingling (Figure 3H). It seems implausible that these thermal modulations reflect variations in some top-down factor such as meaningfulness or interpretability of tactile stimuli. Rather, the temperature dependent modulation may, instead, be explained by bottom-up factors, such as the increase of RA firing at higher temperatures [49,50] and the increase in SA channel activity at lower temperatures [24,58,59].

Another alternative explanation is based on the effective stimulation at the receptors themselves. Recent studies showed that action potentials are accompanied by mechanical deformations of the cell surface [60,61], as well as mechanical waves propagating throughout the axonal surface [62]. Therefore, one possibility is that sustained pressure might have changed the neural response of RA mechanoreceptors or their afferent fibres to sanshool, as a secondary consequence of physically deforming their shape. However, the spatial tuning pattern of our effects offer evidence against this hypothesis. In Experiment 2, sustained touch-related tingling inhibition was strongest at the place of the touch stimulation itself, and at other locations on the same lip. However, we also found that the tingling sensation on the upper lip was modulated by touch on the lower lip and vice versa. Since the lips were held apart during the experiment, this modulation cannot readily be explained by the spread of mechanical input across the lips. Moreover, upper and lower lip are innervated by different branches of the trigeminal nerve (V2 and V3, respectively), which would not allow purely peripheral interactions. Finally, the time-course of suppression is inconsistent with a direct effect of sustained pressure on the RA receptor or itself, or its afferent. In Experiment 4, tingling levels were strongly suppressed immediately after static touch was applied, but then recovered significantly over the subsequent 10 s (Figure 3E). A direct mechanical effect on the RA receptor should presumably remain constant as long as sustained touch lasts. In contrast, the modest recovery of tingle with continuing pressure is consistent with a neural, as opposed to mechanical account, based on the adaptation of SA afferent firing rates.

Our study presents a series of limitations, which should be studied in more detail in the future. First, in our study, channels are defined *perceptually*, and their identification with specific peripheral receptors and afferent fibres can only be putative. Although the physiological characteristics of sanshool are well studied in animal research [17–20], the physiological profile of peripheral mechanoreceptive activation induced by sanshool in humans has not yet been investigated directly, and is known only by psychophysical proxy measures. Future studies could potentially record single peripheral afferents from the human skin microneurographically, and identify the response of different fibre classes to sanshool applied to their respective receptive fields.

Second, the perceptual characteristics of sanshool tingling should be studied in more detail. In our study, we focus on the feature of flutter-level vibration, but other aspects of the sensation remain to be systematically investigated. For example, in a previous study we have shown that sanshool produces tingling at a frequency of 50 Hz [21] and impairs detection of mechanical vibrations at 30 Hz but not higher (240 Hz) or lower (1 Hz) frequencies [22]. The present study extends knowledge of sanshool’s sensory properties by confirming that the perceived intensity of sanshool tingling is dose-dependent (Experiment 3) [32].

Third, the duration of static touch varied widely across our experiments (10 s in Experiment 1, 2 and 4, 1 s in Experiment 3, and 4 s in experiment 6). We varied the duration of static touch because our experiments required different numbers of tactile stimulation trials, which had all to be completed within the typical duration of tingling that follows a single application of sanshool (~40 minutes). Despite the varying tactile durations, we consistently found suppression of tingling sensation, suggesting a rather general effect.

Finally, some of our experiments involved manual delivery of tactile stimuli. These cannot provide precise control over contact force. Given that RA mechanoreceptors are exquisitely sensitive to dynamic changes in contact force, our Experiments 1 and 2 may have included uncontrolled micromovements that activated RA channels. Nevertheless, the precisely-controlled mechanical stimuli of Experiments 3 and 4, which should have drastically reduced micromovements, also produced a strong attenuation of tingling, several seconds after touch onset. These results suggest that the tactile attenuation of tingling is likely to be mediated by an SA rather than by an RA channel activated by unintended micromovements.

At what level in the CNS, then, would putative RA and SA channels interact? Either cortical or sub-cortical interactions are possible. Several circuit mechanisms of presynaptic inhibition have recently been described [63]. In the mouse spinal cord different types of interneurons in the dorsal horn receive inputs from multiple types of low threshold mechanoreceptor (LTMR) al?erents (including RA and SA channels). Since the inhibition of pain by touch (SA channel) is thought to occur at the dorsal horn [6,64,65], the mechanism of analogous inhibition of RA activity by SA input might also be implemented sub-cortically, e.g., spinally or in trigeminal nuclei for somatic or orofacial stimuli respectively. Within somatosensory cortex, neurons in each frequency channel were originally thought to be organised in discrete functional columns[10]. However, many single neurons in area 3b/1 show hybrid activity profiles responding to both RA (transient) and SA (sustained activity) mechanical input [11,15,66]. Therefore, interactions between sub-modalities may occur prior to somatosensory cortex [43].

What might be the functional relevance of a putative SA-RA mechanoreceptor channel interaction? A few studies have previously investigated whether vibration perception is affected by the indentation of the vibrotactile stimulator [67,68]. For example, Lowenthal and colleagues [67] found that detection thresholds for vibratory stimuli are significantly lower at higher contact force levels. However, although seemingly inconsistent with our finding that pressure inhibits vibration perception, Lowenthal’s results may be due to physical interactions between the mechanical stimuli, rather than neural interactions between the resulting signals. More generally, studies with complex mechanical stimuli cannot readily rule out the possibility that apparent interactions between different frequency-tuned tactile channels are in fact due to nonlinear mechanical interactions in the periphery, which influence the effective stimulation at the receptor. In contrast, by using sanshool as a chemical gateway for cutaneous receptors, we were able to reliably deliver tingling sensations in absence of any mechanical confounds (e.g. the pressure exerted by the probe of the vibrotactile stimulator).

To our knowledge, the current study is the first to suggest an inhibitory effect of SA on RA signalling. However, previous reports of an effect in the *reverse* direction, from RA to SA signalling, offer important clues to possible function of such interactions. Bensmaia and colleagues [69,70] showed that increasing the ratio of RA firing to SA firing impaired grating detection performance: RA input interfered with perception of fine spatial structure carried by SA. We therefore speculate that the tactile system contains mechanisms to inhibit RA channel input, to prevent masking by RA-mediated noise, and in order to maintain the robustness and stability of tactile perception. For example, when any tactile contact occurs, mechanical waves [16,71] travel through the skin, and deeper tissues. Interestingly, RA-range frequencies travel over considerable distances. We speculate that SA-induced suppression of RA firing, as reported here, could play an important role in limiting the perceptual impact of these complex mechanical interactions. Lateral inhibition between neurons with adjacent receptive fields is a pervasive feature of sensory spatial representation, serving to increase spatial acuity [72,73]. Lateral inhibition occurs also for non-spatial sensory systems, such as olfaction, where it again serves to enhance perceptual resolution. Our findings are consistent with a functional hypothesis that inhibition of one frequency channel by another frequency channel functions analogously to the enhancement of spatial acuity provided by lateral inhibition. SA-mediated suppression of RA activation during normal touch may serve as a lowpass filter mechanism, allowing reliable perception of tactile events at sensorimotor timescales.

## Data availability

All the data used for the statistical inferences of the paper are available on the Supplementary Material.

## Acknowledgements

This research was supported by an MRC project grant MR/M013901/1 to PH. AC and PH were further supported by a donation by Dr Shamil Chandaria to the Institute of Philosophy, School of Advanced Study, University of London. NH is supported by Japan Society for the Promotion of Science (Kakenhi 26119535, 18H01106) and ERATO (JPMJER1801). YH is supported by The Great Britain Sasakawa Foundation. We are grateful to Indena SPA (Milan, Italy) for providing ZANTHALENE used for the study.

## Author Contribution

AC, NH and PH conceived the study. NH and PH designed, and YH and NH collected and analysed the data for Experiments 1 and 2. AC and PH designed, and AC collected and analysed the data for Experiments 3-6. AC, NH, and PH wrote the paper. All the authors reviewed the paper.

