## Supplementary Material for "Touch inhibits touch: sanshool-induced paradoxical tingling reveals perceptual interference between somatosensory submodalities"

**Supplementary Methods**

***Experiment 1***

We tested whether the intensity of tingling sensation induced by sanshool is modulated by the application of sustained touch.

The target sample size for this experiment (n = 10) was defined a priori, based on our previous studies using sanshool (Hagura et al., 2013; Kuroki et al., 2016).

***Experiment 2***

In Experiment 1, the location where the participants rated the tingling sensation was fixed to one location. In Experiment 2, we confirmed that the result of Experiment 1 is not due to sustained attention to a fixed location of the lips, by varying the location of touch, as well as the location the participant rated the tingling intensity.

The target sample size for this experiment (n = 10) was defined a priori, based on our previous studies using sanshool (Hagura et al., 2013; Kuroki et al., 2016).

***Experiment 3***

In Experiment 3, we tested whether the inhibition of tingling sensation is modulated by the strength of the sustained pressure.

The target sample size for this experiment (n = 8) was defined in a priori, referring to our previous studies using sanshool (Hagura et al., 2013; Kuroki et al., 2016). Nine participants were originally recruited. However, one participant could not follow the instruction of keeping the lips apart (see below), and was therefore excluded from the analyses.

***Experiment 4***

In Experiment 4, we quantified the inhibition of tingling by the sustained pressure by means of amplitude of RA range vibration, and we investigated the temporal dynamics of SA1-RA interaction.

The target sample size was calculated a priori based on the power analysis using data from Experiment 3, aiming to obtain sufficient target power level (alpha = 0.05; power = 0.95; η_p_^2^ = 0.6; estimated sample: n = 12). Eighteen participants originally volunteered for this study. However, two participants could not keep their lips apart, and another two could not perceive strong tingling sensation by sanshool. These participants withdrew from the study, and therefore are not included in the final sample size (n = 14).

In the initial inspection of the data, we found that the distribution of the amplitude data was significantly deviated from the normal distribution (see Supplementary Table S1). Therefore, the statistical analysis was conducted by log-transforming the data. However, to maintain the data in interpretable scale, we report the means and the standard errors in the original units (μm).

***Experiment 5***

Experiment 5 aimed to investigate the contribution of C-nociceptive fibers to the tingling perception induced by Sanshool.

Given the design implemented, participants were only included if they satisfied all of the following conditions: 1) clear sensation of tingle evoked by sanshool; 2) clear nociceptive sensation evoked by noxious electrocutaneous stimulation; 3) effective lidocaine block of nociceptive afference, as measured by reduced pain sensation/increased pain threshold. Of the 14 participants originally recruited, one was excluded during the pre-test evaluation of sanshool-induced tingling, because administration of sanshool did not elicit any tingling on the lips after 10 minutes from application; another was excluded during the pre-test evaluation of pain thresholds, because electrocutaneous stimulation within the safe intensity range did not elicit any painful sensation. Four further participants were excluded in the post-test evaluation of pain, because they did not show any numerical increase in pain threshold after lidocaine administration.

Given the Bayesian analyses approach used in this study, and the use of painful stimuli, data collection was interrupted after reaching the minimal significant result supporting either hypotheses.

***Experiment 6***

Experiment 6 took place in the context of a science exhibition at the Tate Modern (London). A total of 64 attendees took part in the event. However, only the data from participants respecting all the following inclusion criteria were used for further analysis: 1) being between 18 and 65 years old; 2) not wearing any lipstick, lip gloss, or lip balm; 3) feeling a clear tingling sensation evoked by sanshool (i.e. baseline rating ≥ 3); 4) completing all the trials of the experiment. This left a final sample size of 51 participants.

**Supplementary Results**

***Experiment 1***

To investigate the spatial gradient of the tingle inhibition by probe touch, we directly compared the tingle ratings across different touch locations on the upper and lower lip. 2 (lip; target position same or different to the probe touch) x 3 (position; three horizontal locations on the lip) ANOVA revealed significant main effect both for the factor of the lip (F(1,9) = 20.33, *p* = 0.001, η_p_^2^ = 0.693) and the position of the probe touch (F(2,18) = 8.43, *p* = 0.003, η_p_^2^ = 0.484). An interaction effect was also significant (F(2,18) = 5.28, *p* = 0.016, η_p_^2^ = 0.370). To investigate the nature of the interaction effect, ratings were also analysed within each lip. One-way ANOVA on ratings of lower lip touch (positions 5, 6 and 7) revealed significant main effect between the three different probe positions (F(2,18) = 10.89, *p* < 0.001, η_p_^2^ = 0.547). Planned comparisons showed that tingle intensity was reduced more strongly by pressure at position 6 (target position) than at position 5 (t(9) = 3.05, *p* = 0.014, dz = 0.96) and position 7 (t(9) = 4.32, *p* = 0.002, dz = 1.37) . For the upper lip (positions 2, 3 and 4), although the average rating was numerically lower for the middle position (position 3; 74.2% of baseline intensity) it was not significantly different from the adjacent two locations (position 2; 81.2%, position 4; 79.2%) (F(2,18) = 10.868, *p* = 0.55, η_p_^2^ = 0.065).

**Supplementary Figures**

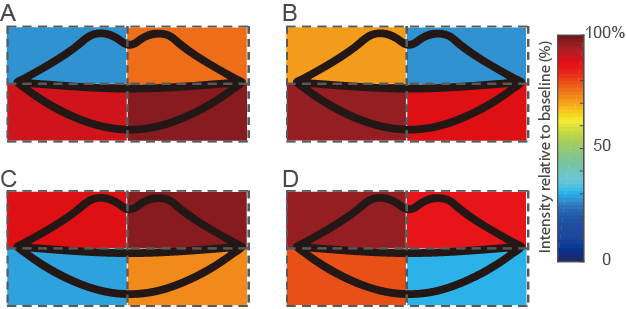

**Supplementary Figure S1:** Reduction of sanshool tingling intensity on the lips in Experiment 2. Sustained force was applied to different quadrants of the lip (A: Left upper, B: Right upper, C: Left lower, D: Right lower), while participants experienced sanshool induced tingling in all of the lip locations. The tingling intensity was robustly inhibited at the quadrant where the sustained touch was applied, with less suppression in other, untouched quadrants.

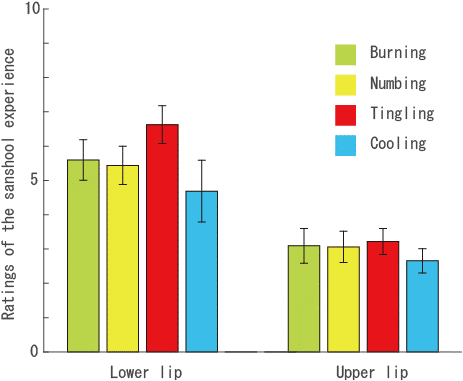

**Supplementary Figure S2:** Manipulation check to confirm effect of sanshool solution strength on sensations in Experiment 3. 80% solution of sanshool was applied to the lower lip, whereas 20% solution of sanshool was applied to the upper lip, aiming to induce stronger intensity of tingling experience on the lower lip compared to the upper lip. Participants reported the sanshool induced tingling experience using numerical ratings (0 to 10, 0: no sensation, 10: strongest imaginable sensation). They were also asked to report other sensations which have been associated with sanshool; burning, numbing and cooling (Hagura et al., 2013). For all the descriptors, the lower lip had higher ratings. Tingling sensation, which is our main interest, was significantly stronger for the lower lip than the upper lip (t(7) = 6.94, p < 0.001, dz = 2.45). This shows that our manipulation to induce different levels of tingling intensity using different solution strengths on upper and lower lip was successful. Error bars indicate standard error of the mean across participants (n = 8).

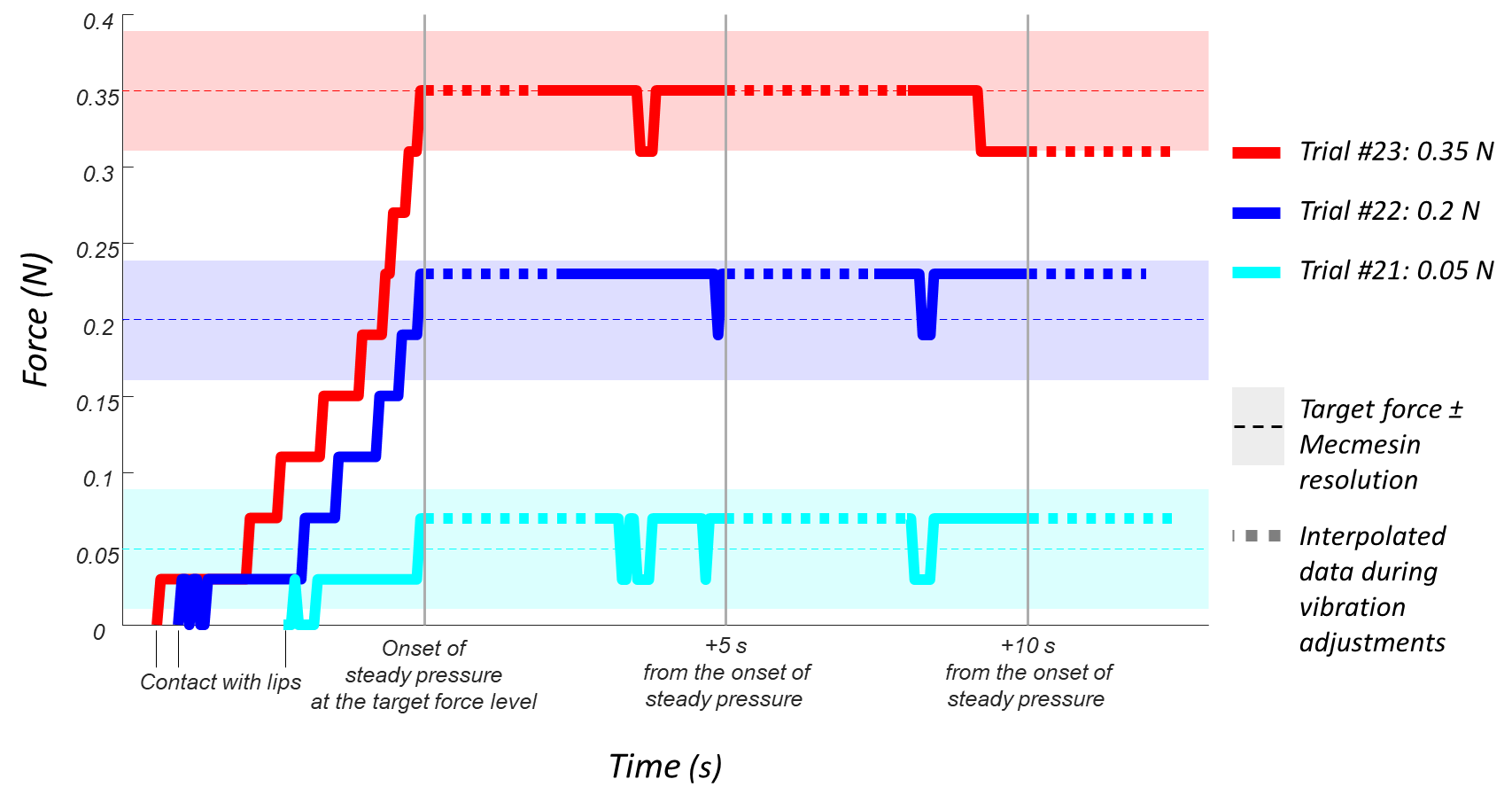

**Supplementary Figure S3.** Force traces from three trials of a representative participant in Experiment 4. The continuous coloured lines represent the reading of the Mecmesin force sensor from the moment of contact between the probe and the lips until the end of the trial. Three trials are shown, with force target levels of 0.35 N (red trace), 0.2 N (blue trace), and 0.05 N (cyan trace). The dashed portions of each trace represent interpolated data for the period in which participants adjusted a mechanical vibration to the upper lip to match the intensity of sanshool-evoked tingle on the lower lip. During this adjustment period the force could not be recorded. The duration of each adjustment depended on the participant themselves (see Methods and Supplementary Video S1). Thin dashed horizontal lines indicate the target force for each trial. Surrounding coloured bands indicate the resolution of the Mecmesin PFI-200N (± 0.04 N) according to the manufacturer’s technical specification.

**Supplementary Tables**

**Supplementary Table S1.** Table of means of Experiment 1.

|  | **Rating of tingling on the target position 6 when the position touched is:** | | | | | | | |
| --- | --- | --- | --- | --- | --- | --- | --- | --- |
| **Subj #** | **1** | **2** | **3** | **4** | **5** | **6** | **7** | **8** |
| **1** | 5.50 | 7.17 | 5.42 | 5.33 | 3.17 | 0.83 | 4.50 | 5.17 |
| **2** | 10.00 | 7.50 | 7.92 | 10.33 | 9.00 | 9.58 | 8.50 | 8.67 |
| **3** | 7.33 | 10.50 | 4.75 | 7.67 | 7.33 | 1.75 | 3.83 | 7.00 |
| **4** | 7.83 | 8.33 | 8.83 | 7.50 | 5.83 | 0.33 | 5.50 | 8.50 |
| **5** | 14.50 | 7.83 | 10.58 | 9.83 | 5.33 | 7.42 | 9.00 | 14.33 |
| **6** | 8.33 | 10.00 | 8.17 | 6.67 | 4.83 | 0.00 | 5.00 | 9.50 |
| **7** | 9.67 | 10.83 | 11.75 | 11.67 | 5.67 | 0.92 | 6.17 | 10.83 |
| **8** | 4.50 | 3.67 | 1.50 | 4.17 | 2.50 | 0.17 | 4.67 | 5.33 |
| **9** | 10.33 | 10.67 | 11.33 | 10.67 | 11.67 | 3.67 | 11.67 | 11.00 |
| **10** | 8.17 | 4.67 | 3.92 | 5.33 | 0.17 | 0.00 | 1.33 | 11.50 |

**Supplementary Table S2.** Table of means of Experiment 2.

|  | **Rating of tingling in each quadrant when position 1 is touched** | | | | **Rating of tingling in each quadrant when position 2 is touched** | | | | **Rating of tingling in each quadrant when position 3 is touched** | | | | **Rating of tingling in each quadrant when position 4 is touched** | | | |
| --- | --- | --- | --- | --- | --- | --- | --- | --- | --- | --- | --- | --- | --- | --- | --- | --- |
| **S#** | **1** | **2** | **3** | **4** | **1** | **2** | **3** | **4** | **1** | **2** | **3** | **4** | **1** | **2** | **3** | **4** |
| **1** | 0.3 | 5.3 | 4.3 | 6.5 | 4.0 | 0.0 | 7.2 | 3.7 | 4.5 | 7.2 | 0.5 | 3.5 | 7.0 | 3.2 | 4.7 | 0.3 |
| **2** | 1.3 | 4.7 | 9.0 | 9.5 | 3.7 | 1.5 | 9.8 | 8.5 | 6.8 | 9.5 | 1.2 | 3.8 | 9.3 | 6.5 | 7.2 | 1.5 |
| **3** | 10.0 | 10.3 | 10.3 | 11.3 | 10.5 | 9.8 | 11.2 | 11.0 | 10.7 | 10.0 | 10.0 | 11.7 | 10.7 | 11.7 | 10.8 | 10.3 |
| **4** | 0.5 | 10.2 | 9.0 | 9.7 | 9.5 | 0.3 | 9.3 | 9.8 | 10.0 | 10.0 | 0.8 | 9.0 | 9.8 | 10.2 | 9.0 | 0.7 |
| **5** | 6.5 | 10.0 | 9.7 | 10.0 | 9.0 | 6.2 | 9.0 | 9.2 | 10.0 | 10.0 | 7.3 | 10.0 | 10.0 | 10.0 | 10.0 | 7.0 |
| **6** | 0.0 | 8.0 | 9.7 | 10.0 | 7.3 | 0.0 | 8.7 | 7.3 | 10.0 | 9.5 | 0.0 | 6.3 | 10.0 | 9.7 | 8.0 | 0.0 |
| **7** | 0.8 | 6.2 | 4.3 | 6.7 | 4.3 | 0.7 | 5.2 | 5.8 | 2.0 | 5.8 | 0.7 | 3.3 | 4.7 | 4.8 | 4.7 | 1.3 |
| **8** | 0.3 | 3.7 | 8.2 | 10.5 | 5.7 | 1.3 | 8.3 | 6.3 | 9.2 | 7.3 | 0.8 | 7.5 | 5.7 | 5.7 | 5.8 | 2.0 |
| **9** | 7.2 | 11.0 | 10.3 | 10.3 | 10.3 | 7.0 | 10.7 | 10.7 | 11.8 | 12.2 | 7.3 | 11.0 | 12.0 | 11.3 | 10.3 | 7.0 |
| **10** | 0.0 | 6.2 | 14.8 | 13.6 | 6.2 | 0.4 | 16.4 | 15.4 | 13.6 | 16.2 | 0.0 | 6.6 | 16.2 | 13.6 | 8.0 | 0.0 |

**Supplementary Table S3.** Table of means of Experiment 3.

|  | **Probability (%) of reporting that the tingling on the lower lip is stronger than the tingling on the upper lip when force applied on the lower lip is:** | | | | |
| --- | --- | --- | --- | --- | --- |
| **Subj #** | **0.05 N** | **0.16 N** | **0.28 N** | **0.39 N** | **0.50 N** |
| **1** | 100.0 | 70.0 | 60.0 | 23.3 | 6.7 |
| **2** | 90.0 | 50.0 | 33.3 | 40.0 | 13.3 |
| **3** | 80.0 | 50.0 | 53.3 | 36.7 | 30.0 |
| **4** | 86.7 | 73.3 | 43.3 | 50.0 | 40.0 |
| **5** | 46.7 | 46.7 | 56.7 | 50.0 | 46.7 |
| **6** | 70.0 | 56.7 | 60.0 | 40.0 | 26.7 |
| **7** | 90.0 | 70.0 | 40.0 | 50.0 | 23.3 |
| **8** | 60.0 | 50.0 | 56.7 | 53.3 | 56.7 |

**Supplementary Table S4.** Table of means of Experiment 4.

|  | **Baseline** | | | **Pressure onset**  **(0 s)** | | | **+5 s from pressure onset** | | | **+10 s from pressure onset** | | |
| --- | --- | --- | --- | --- | --- | --- | --- | --- | --- | --- | --- | --- |
| **Subj #** | **0.05 N** | **0.2**  **N** | **0.35**  **N** | **0.05 N** | **0.2**  **N** | **0.35**  **N** | **0.05 N** | **0.2**  **N** | **0.35**  **N** | **0.05 N** | **0.2**  **N** | **0.35**  **N** |
| **1** | 11.5 | 12.7 | 11.5 | 10.2 | 11.5 | 11.1 | 9.6 | 10.9 | 11.0 | 10.4 | 10.9 | 10.2 |
| **2** | 29.0 | 31.0 | 28.3 | 17.9 | 15.3 | 14.5 | 21.3 | 18.0 | 16.5 | 20.3 | 21.8 | 19.8 |
| **3** | 11.3 | 11.3 | 11.8 | 6.0 | 5.8 | 5.2 | 6.5 | 5.5 | 6.1 | 7.0 | 6.4 | 6.1 |
| **4** | 7.5 | 11.2 | 8.4 | 8.1 | 9.3 | 6.8 | 8.0 | 12.8 | 6.9 | 7.0 | 7.4 | 8.9 |
| **5** | 14.7 | 17.1 | 22.2 | 9.8 | 6.6 | 5.3 | 9.1 | 6.8 | 4.1 | 9.8 | 6.5 | 3.9 |
| **6** | 17.9 | 18.8 | 17.9 | 9.3 | 10.6 | 7.8 | 9.3 | 9.8 | 7.5 | 11.3 | 8.2 | 7.4 |
| **7** | 18.8 | 20.3 | 21.7 | 7.6 | 4.9 | 4.8 | 7.7 | 5.5 | 6.4 | 9.6 | 7.5 | 5.8 |
| **8** | 14.0 | 14.2 | 12.9 | 8.2 | 11.7 | 10.3 | 7.6 | 14.2 | 13.8 | 8.2 | 16.6 | 16.7 |
| **9** | 13.6 | 13.4 | 11.7 | 7.4 | 7.1 | 6.3 | 7.3 | 7.9 | 7.9 | 7.2 | 7.7 | 7.2 |
| **10** | 12.6 | 12.6 | 12.0 | 5.3 | 4.0 | 3.1 | 6.1 | 4.1 | 3.0 | 6.1 | 4.5 | 3.2 |
| **11** | 12.6 | 11.3 | 10.6 | 10.6 | 8.7 | 4.7 | 10.5 | 10.2 | 6.1 | 10.7 | 8.2 | 6.4 |
| **12** | 7.1 | 7.6 | 6.9 | 6.0 | 5.9 | 6.1 | 6.6 | 6.6 | 5.8 | 5.8 | 5.9 | 5.8 |
| **13** | 13.2 | 12.9 | 11.2 | 5.7 | 6.4 | 6.5 | 6.5 | 6.3 | 6.3 | 7.2 | 7.1 | 6.8 |
| **14** | 7.3 | 6.2 | 7.0 | 2.5 | 2.2 | 2.1 | 2.7 | 2.2 | 2.1 | 2.7 | 2.4 | 2.1 |

**Supplementary Table S5.** Table of individual data from Experiment 5.

| **Subj #** | **Pain threshold PRE (mA)** | **Pain threshold POST (mA)** | **Tingling rating PRE (0-10)** | **Tingling rating POST (0-10)** |
| --- | --- | --- | --- | --- |
| 1 | 0.8 | 0.9 | 7.0 | 7.0 |
| 2 | 0.7 | 0.9 | 7.0 | 7.0 |
| 3 | 1.1 | 1.6 | 8.0 | 5.0 |
| 4 | 0.6 | 1.0 | 4.0 | 4.5 |
| 5 | 0.8 | 0.9 | 6.0 | 7.0 |
| 6 | 0.7 | 1.1 | 6.0 | 6.0 |
| 7 | 0.8 | 1.0 | 3.0 | 3.0 |
| 8 | 1.4 | 1.5 | 5.0 | 7.0 |

**Supplementary Table S6.** Table of means of Experiment 6.

|  | | **Rating of tingling (VAS) when the tactile stimulus is at:** | | |
| --- | --- | --- | --- | --- |
| **Subj #** | **Baseline Tingle**  **(0-10)** | **21°C** | **33°C** | **41°C** |
| **1** | 4 | 0.0 | 0.1 | 5.2 |
| **2** | 3 | 6.7 | 3.5 | 4.9 |
| **3** | 5 | 2.0 | 3.6 | 6.1 |
| **4** | 3 | 3.0 | 4.3 | 7.7 |
| **5** | 4 | 0.3 | 1.8 | 2.0 |
| **6** | 7 | 1.0 | 1.8 | 4.5 |
| **7** | 8 | 3.1 | 6.1 | 7.2 |
| **8** | 6 | 1.2 | 0.7 | 2.2 |
| **9** | 5 | 3.2 | 5.0 | 8.6 |
| **10** | 6 | 2.6 | 4.8 | 7.5 |
| **11** | 5 | 0.6 | 0.9 | 0.0 |
| **12** | 10 | 1.1 | 4.6 | 5.3 |
| **13** | 4 | 1.2 | 2.0 | 2.2 |
| **14** | 3 | 1.5 | 6.8 | 9.7 |
| **15** | 10 | 0.0 | 0.2 | 0.2 |
| **16** | 6 | 4.9 | 5.9 | 7.6 |
| **17** | 6 | 1.5 | 2.0 | 7.3 |
| **18** | 8 | 0.9 | 3.1 | 3.9 |
| **19** | 4 | 4.7 | 2.4 | 6.5 |
| **20** | 6 | 0.2 | 1.3 | 1.3 |
| **21** | 3 | 0.7 | 1.3 | 3.5 |
| **22** | 7 | 0.1 | 1.7 | 2.4 |
| **23** | 8 | 1.2 | 4.3 | 2.0 |
| **24** | 5 | 1.6 | 2.2 | 4.9 |
| **25** | 7 | 1.8 | 0.5 | 1.3 |
| **26** | 6 | 4.1 | 4.9 | 6.0 |
| **27** | 5 | 0.9 | 4.3 | 6.4 |
| **28** | 8 | 1.2 | 3.0 | 8.2 |
| **29** | 10 | 0.1 | 2.0 | 6.4 |
| **30** | 8 | 3.9 | 2.0 | 3.9 |
| **31** | 4 | 0.7 | 4.8 | 8.2 |
| **32** | 8 | 0.2 | 3.1 | 4.4 |
| **33** | 4 | 3.4 | 5.2 | 7.3 |
| **34** | 8 | 1.5 | 0.9 | 0.4 |
| **35** | 7 | 8.1 | 4.0 | 3.7 |
| **36** | 6 | 2.1 | 3.2 | 4.9 |
| **37** | 7 | 0.8 | 0.9 | 4.1 |
| **38** | 6 | 4.3 | 2.4 | 5.1 |
| **39** | 9 | 1.4 | 3.7 | 8.4 |
| **40** | 8 | 3.7 | 3.1 | 2.7 |
| **41** | 6 | 0.8 | 2.2 | 1.6 |
| **42** | 4 | 0.6 | 0.1 | 4.0 |
| **43** | 7 | 1.5 | 4.9 | 3.2 |
| **44** | 8 | 1.7 | 2.0 | 2.7 |
| **45** | 6 | 0.3 | 0.0 | 0.2 |
| **46** | 6 | 1.8 | 4.0 | 6.1 |
| **47** | 6 | 1.1 | 4.5 | 6.5 |
| **48** | 5 | 0.1 | 1.8 | 7.7 |
| **49** | 6 | 2.1 | 1.5 | 1.0 |
| **50** | 6 | 1.9 | 2.5 | 6.6 |
| **51** | 3 | 0.7 | 0.6 | 4.9 |

**Supplementary Table S7. Normality test for Experiment 4.**

|  | **Shapiro-Wilk test** | | |
| --- | --- | --- | --- |
| **Condition** | **Statistic** | **df** | **p-value** |
| **0.05_Baseline** | 0.860 | 14 | 0.031 |
| **0.05_0s** | 0.887 | 14 | 0.073 |
| **0.05_5s** | 0.738 | 14 | 0.001 |
| **0.05_10s** | 0.843 | 14 | 0.018 |
| **0.2_Baseline** | 0.866 | 14 | 0.037 |
| **0.2_0s** | 0.970 | 14 | 0.877 |
| **0.2_5s** | 0.957 | 14 | 0.677 |
| **0.2_10s** | 0.799 | 14 | 0.005 |
| **0.35_Baseline** | 0.861 | 14 | 0.031 |
| **0.35_0s** | 0.919 | 14 | 0.214 |
| **0.35_5s** | 0.889 | 14 | 0.078 |
| **0.35_10s** | 0.833 | 14 | 0.013 |

**Supplementary Video S1**

A video showing the setup in Experiment 3 and 4, and an example trial from Experiment 4 can be found at <https://tinyurl.com/yyuoecqd>.
